# Automated recognition of functional compound-protein relationships in literature

**DOI:** 10.1101/718205

**Authors:** Kersten Döring, Ammar Qaseem, Kiran K Telukunta, Michael Becer, Philippe Thomas, Stefan Günther

**Author notes:** These authors contributed equally to this work (Kersten Döring, Ammar Qaseem).

## Abstract

**Motivation:** Much effort has been invested in the identification of protein-protein interactions using text mining and machine learning methods. The extraction of functional relationships between chemical compounds and proteins from literature has received much less attention, and no ready-to-use open-source software is so far available for this task.

**Method:** We created a new benchmark dataset of 2,753 sentences from abstracts containing annotations of proteins, small molecules, and their relationships. Two kernel methods were applied to classify these relationships as functional or non-functional, named shallow linguistic and all-paths graph kernel. Furthermore, the benefit of interaction verbs in sentences was evaluated.

**Results:** The cross-validation of the all-paths graph kernel (AUC value: 84.2%, F1 score: 81.8%) shows slightly better results than the shallow linguistic kernel (AUC value: 81.6%, F1 score: 79.7%) on our benchmark dataset. Both models achieve state-of-the-art performance in the research area of relation extraction. Furthermore, the combination of shallow linguistic and all-paths graph kernel could further increase the overall performance. We used each of the two kernels to identify functional relationships in all PubMed abstracts (28 million) and provide the results, including recorded processing time.

**Availability:** The software for the tested kernels, the benchmark, the processed 28 million PubMed abstracts, all evaluation scripts, as well as the scripts for processing the complete PubMed database are freely available at https://github.com/KerstenDoering/CPI-Pipeline.

**Author summary:** Text mining aims at organizing large sets of unstructured text data to provide efficient information extraction. Particularly in the area of drug discovery, the knowledge about small molecules and their interactions with proteins is of crucial importance to understand the drug effects on cells, tissues, and organisms. This data is normally hidden in written articles, which are published in journals with a focus on life sciences. In this publication, we show how text mining methods can be used to extract data about functional interactions between small molecules and proteins from texts. We created a new dataset with annotated sentences of scientific abstracts for the purpose of training two diverse machine learning methods (kernels), and successfully classified compound-protein pairs as functional and non-functional relations, i.e. no interactions. Our newly developed benchmark dataset and the pipeline for information extraction are freely available for download. Furthermore, we show that the software can be easily up-scaled to process large datasets by applying the approach to 28 million abstracts.

## Introduction

Interactions of biomolecules are substantial for most cellular processes, involving metabolism, signaling, regulation, and proliferation [1]. Small molecules (compounds) can serve as substrates by interacting with enzymes, as signal mediators by binding to receptor proteins, or as drugs by interacting with specific target proteins [2].

Detailed information about compound-protein interactions is provided in several databases. ChEMBL annotates binding affinity and activity data of small molecules derived from diverse experiments [3]. PDBbind describes binding kinetics of ligands that have been co-crystallized with proteins [4]. DrumPID focuses on drugs and their addressed molecular networks including main and side-targets [5]. DrugBank, SuperTarget, and Matador host information mainly on FDA approved but also experimental drugs and related interacting proteins [6, 7]. However, most of this data was extracted from scientific articles. Given that more than 10,000 new articles are added in the literature database PubMed each week, it is obvious that it requires much effort to extract and annotate these information manually to generate comprehensive datasets. Automatic text mining methods may support this process significantly [1, 8]. Today, only a few approaches exist for this specific task. One of them is the Search Tool Interacting Chemicals (STITCH), developed in its 5th version in 2016, which connects several information sources of compound-protein interactions [2]. This includes experimental data and data derived from text mining methods based on co-occurrences and natural language processing [9, 10]. Similar methods have been applied for developing the STRING 11.0 database, which contains mainly protein-protein interactions [11]. OntoGene is a text mining web service for the detection of proteins, genes, drugs, diseases, chemicals, and their relationships [12]. The identification methods contain rule-based and machine learning approaches, which were successfully applied in the BioCreative challenges, e.g. in the triage task in 2012 [13].

Although STITCH and OntoGene deliver broadly beneficial text mining results, it is difficult to compare their approaches, because no exact statistical measures of their protein-compound interaction prediction methods are reported. Furthermore, no published gold standard corpus of annotated compound-protein interactions could be found to evaluate text mining methods for their detection.

Tikk *et al.* compared 13 kernel methods for protein-protein interaction extraction on several text corpora. Out of these methods, the SL kernel and APG kernel consistently achieved very good results [14]. To detect binary relationships, the APG kernel considers all weighted syntactic relationships in a sentence based on a dependency graph structure. In contrast, the SL kernel considers only surface tokens before, between, and after the candidate interaction partners.

Both kernels have been successfully applied in different domains, including drug-drug interaction extraction [15], the extraction of neuroanatomical connectivity statements [16], and the I2B2 relation extraction challenge [17].

If two biomolecules appear together in a text or a sentence, they are referred to as co-occurring. A comparably high number of such pairs of biomolecules can be used as a prediction method for functional relationship detection, e.g. between proteins or proteins and chemical compounds [9]. This concept can be refined by requiring interaction words such as specific verbs in a sentence [1, 18].

So far, it was unclear whether machine learning outperforms a rather naive baseline relying on co-occurrences (with or without interaction words) for the detection of functional compound-protein relations in texts. Especially for the identification of newly described interactions in texts, that were not described in any other data source, the annotation of a functional relation is challenging.

In this publication, we evaluated the usability of two diverse state-of-the-art machine learning kernels for the detection of functional and non-functional compound-protein relationships in texts, independent of additional descriptors, such as the frequency of co-occurrences of specific pairs. To achieve the goal of benchmarking, we annotated a corpus of protein and compound names in 2,753 sentences and manually classified their relations as functional and non-functional, i.e. no interaction. Furthermore, the kernels were applied to the large-scale text dataset of PubMed with 28 million abstracts. The approaches have been implemented in an easy-to-use open source software available via GitHub.

Both kernels achieved a better performance on the benchmark dataset than simply using the concept of co-occurrences. These findings imply that a relatively high classification threshold can be used to automatically identify and extract functional compound-protein interactions from publicly available literature with high precision.

## Dataset and methods

### Generation of the benchmark dataset

Chemical compounds are referred to as small molecules up to a molecular weight of about 1,000 Da, for which a synonym and a related ID are contained in the PubChem database [19]. Similarly, genes and proteins must have UniProt synonyms and were assigned to related UniProt IDs [20].

PubChem synonyms were automatically annotated with the approach described in the manuscripts about the web services CIL [21] and prolific [22], by applying the rules described by Hettne *et al.* [23]. Proteins were annotated using the web service Whatizit [24]. Synonyms that were assigned wrongly by the automatic named entity recognition approach were manually removed.

The complete compound-protein-interaction benchmark dataset (CPI-DS) was generated from the first 40,000 abstracts of all PubMed articles published in 2009, using PubMedPortable [25].

All pairs of proteins and compounds co-occurring in a sentence are considered as potential functionally related or putative positive instances. Pairs with no functional relation were subsequently annotated as negative instances. If a named entity exists as a long-form synonym and an abbreviated form in brackets, both terms are considered as individual entities. The result is a corpus of 2,753 sentences containing at least one compound and protein name (CPI-pair).

For further manual annotation, all sentences were transferred to an HTML form. Verbs that belong to a list of defined interaction verbs, defined by Senger *et al.* [22] and which are enclosed by a protein-compound pair, were annotated, too. All detected CPI-pairs were manually classified as functionally related (positive instances) and non-functionally related CPI-pairs (negative instances).

### Interaction verbs

To analyze how much specific verbs, enclosed by a compound and protein name, affect the precision of functional relationships, we further differentiated between sentences with or without this structure. Fig 1 shows detailed examples.

**Fig 1.**
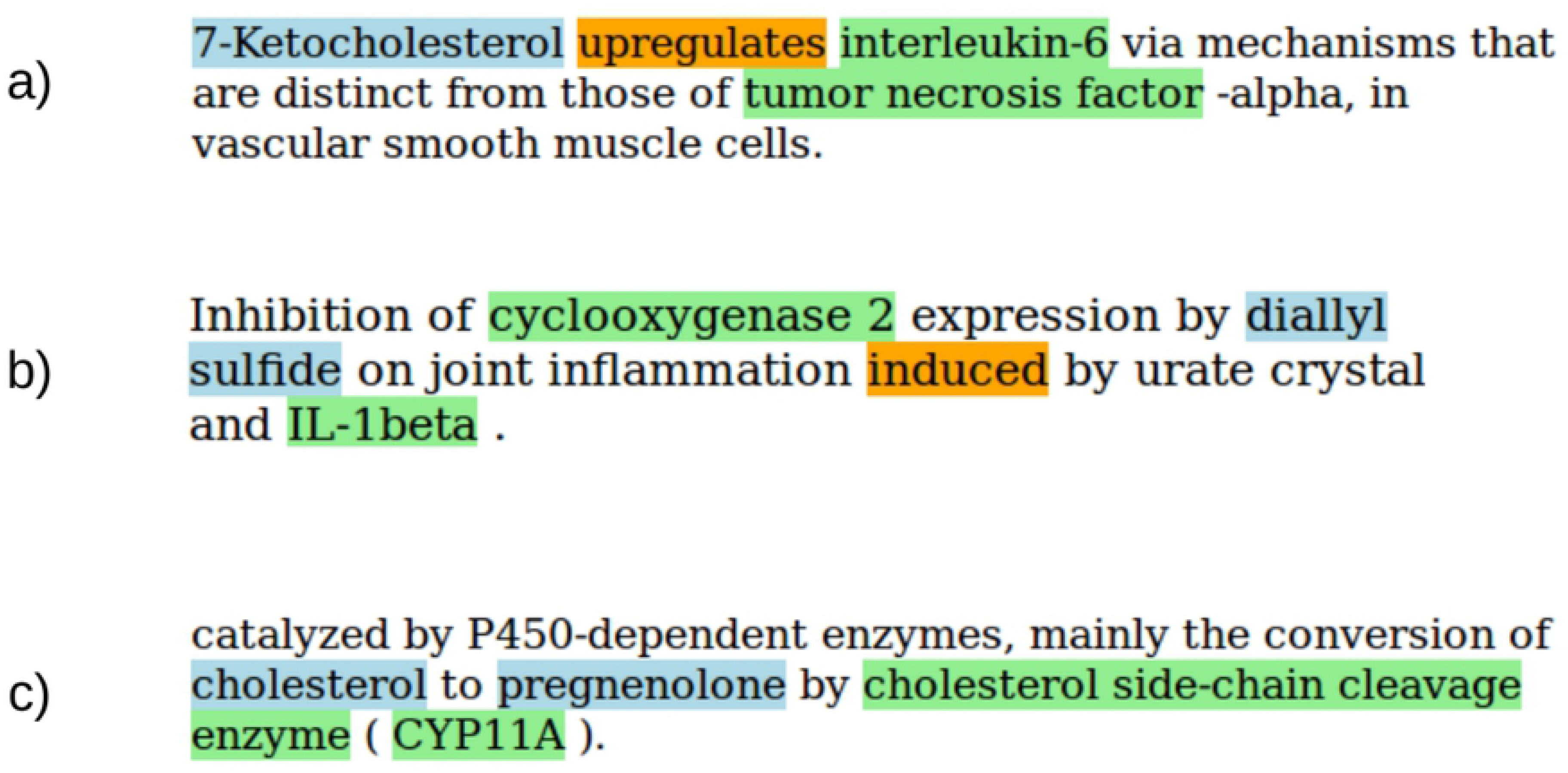
a) **Direct functional relation with interaction verb**. The orange-coloured verb is enclosed by the compound 7-ketocholesterol, shown in blue, and the protein interleukin-6, shown in green. The pair was annotated as functional. b) **Indirect functional relation with interaction verb**. Diallyl sulfide is influencing cyclooxygenase 2 indirectly by inhibiting its expression. The pair was annotated as functional. The compound diallyl sulfide and the protein IL-1beta enclose an interaction verb, but do not describe a functional relation. c) **Direct functional relation without interaction verb**. The molecule cholesterol is metabolised to pregnenolone by CYP11A. This is indicated by the word conversion. The pair was annotated as functional.

### Kernel methods

#### Shallow linguistic kernel

Giuliano *et al.* developed this kernel to perform relation extraction from biomedical literature [26]. It is defined as the sum of a global and local context kernel. These customized kernels were implemented with the LIBSVM framework [27]. Tikk *et al.* adapted the LIBSVM package to compute the distance to the hyperplane, which allowed us to calculate an area under the curve value.

The global context kernel works on unsorted patterns of words up to a length of *n*=3. These n-grams are implemented using the bag-of-words approach. The method counts the number of occurrences of every word in a sentence including punctuation, but excludes the candidate entities. The patterns are computed regarding the phrase structures before-between, between, and between-after the considered entities.

The local context kernel considers tokens with their part-of-speech tags, capitalisation, punctuation, and numerals [1, 26]. The left and right ordered word neighborhoods up to window size of *w*=3 are considered in two separated kernels, which are summed up for each relationship instance.

#### All-paths graph kernel

The APG kernel is based on dependency graph representations of sentences, which are gained from dependency trees [28]. In general, the nodes in the dependency graph are the text tokens in the text (including the part-of-speech tag). The edges represent typed dependencies, showing the syntax of a sentence. The highest emphasis is given to edges which are part of the shortest path connecting the compound-protein pair in question. A graph can be represented in an adjacency matrix. The entries in this matrix determine the weights of the connecting edges. A multiplication of the matrix with itself returns a new matrix with all summed weights of path length two.

All possible paths of all lengths can be calculated by computing the powers of the matrix. Matrix addition of all these matrices results in a final adjacency matrix, which consists of the summed weights of all possible paths [30]. Paths of length zero are removed by subtracting the identity matrix.

All labels are represented as a feature vector. The feature vector is encoded for every vertex, containing the value 1 for labels that are presented within this particular node. This results in a label allocation matrix.

A feature matrix as defined by Gärtner *et al.* sums up all weighted paths with all presented labels [29]. This calculation combines the strength of the connection between two nodes with the encoding of their labels. In general, it can be stated that the dependency weights are higher the shorter their distance to the shortest path between the candidate entities is [1]. The similarity of two feature matrix representations can be computed by summing up the products of all their entries [30].

In the implementation used here [1, 30], the regularized least squares classifier algorithm is applied to classify compound-protein interactions with the APG kernel. This classifier is similar to a standard support vector machine, but the underlying mathematical problem does not need to be solved with quadratic programming [30, 31].

### Analysis of predictive models

#### Baseline of the kernel models

We considered co-occurrences as a simple approach to calculate the baseline in the way that every appearance of a compound and a protein in a sentence is classified as a functional relationship (recall 100%, specificity 0%), taking into account the number of all true functional relationships.

### Benchmark dataset-based analysis

The evaluation was calculated by document-wise 10-fold cross-validation, as an instance-wise cross-validation leads to overoptimistic performance estimates [32]. Each compound-protein pair was classified as functionally related or not related using the previously described kernel methods and resulting in an overall recall, precision, F1 score, and AUC value for a range of kernel parameters.

Subsequently, the kernels were applied solely to sentences which contain an interaction verb and sentences containing no interaction verb. The analyses compare each of the three baselines with the kernel method results.

### Combination of the APG and SL kernel

To analyze if the combination of both kernels yields a higher precision and F1 score than each individual kernel, we combined them by applying a jury decision rule, with the definition that only those relations were classified as functional for which both kernels predicted a functional relation. As we are considering the benchmark dataset, the values for recall, specificity, precision, accuracy, and F1 score could be calculated for the new jury outcome. Based on what is known from the AUC analysis, we can identify the classification threshold for which each of the two kernels reaches the same precision as resulting from the jury decision with default thresholds. By recalculating the above mentioned parameters for each kernel with the new classification threshold, we can compare the single kernel performance with the jury decision outcome.

### Large-scale dataset analysis

We applied both kernels on all PubMed abstracts before 2018, including titles. For the annotation of proteins and small molecules the PubTator web service was applied [33, 34]. The web server annotates genes, proteins, compounds, diseases, and species in uploaded texts. Furthermore, it provides all PubMed abstracts and titles as preprocessed data.

Processing these annotations with PubMedPortable [25] in combination with the CPI pipeline, as explained in the GitHub project documentation, allows for a complete automatic annotation of functional compound-protein relations in texts.

## Results and discussion

### Baseline analysis

Within all sentences, a total number of 5,963 CPI-pairs was curated and separated into 3,496 functionally related (positive instances) and 2,467 non-functionally related CPI-pairs (negative instances). Considering the prediction approach of co-occurrences, this results in a precision (equal to accuracy in this case) of 58.6% and an F1 score of 73.9% (Table 1).

**Table 1.** Analysis of the CPI benchmark dataset.

| DS | #Sent. | #CPIs | #No-CPIs | Total | Rec. | Spec. | Prec. | F<sub>1</sub> |
| --- | --- | --- | --- | --- | --- | --- | --- | --- |
| CPI-DS | 2753 | 3496 | 2467 | 5963 | 100.0 | 0.0 | 58.6 | 73.9 |

Baseline results for precision, recall, and F1 score based on simple co-occurrences. Results are shown in percent (DS - dataset, Sent. - sentences, Rec. - recall, Spec. - specificity, Prec. - precision, F1 - F1 score).

### Shallow Linguistic Kernel

All parameter combinations in the range 1-3 for n-gram and window size of the SL kernel were evaluated. The selection of n-gram 3 and window size 1 shows the best AUC value and the highest precision in comparison to all other models. In general, a lower value of window size leads to a higher precision and a lower recall (Table 2).

**Table 2.** Shallow linguistic kernel results on the datasets CPI-DS.

| n | w | Rec. | Spec. | Prec. | F1 | AUC |
| --- | --- | --- | --- | --- | --- | --- |
| 1 | 1 | 77.3 | 69.0 | 78.1 | 77.7 | 79.2 |
| 1 | 2 | 83.1 | 62.0 | 75.7 | 79.2 | 78.6 |
| 1 | 3 | 87.6 | 52.8 | 72.6 | 79.2 | 78.0 |
| 2 | 1 | 79.2 | 69.9 | 79.1 | 79.1 | 80.9 |
| 2 | 2 | 84.1 | 62.4 | 76.2 | 79.9 | 80.2 |
| 2 | 3 | 87.9 | 55.0 | 73.6 | 80.0 | 79.9 |
| 3 | 1 | 80.3 | 69.7 | 79.2 | 79.7 | 81.6 |
| 3 | 2 | 84.0 | 63.5 | 76.7 | 80.1 | 81.0 |
| 3 | 3 | 87.8 | 56.0 | 74.0 | 80.2 | 80.8 |

The first parameter shows the n-gram value, the second represents the window size. Values in percent: Rec. - recall, Spec. - specificity, Prec. - precision, F1 - F1 score, AUC - area under the curve.

### All-paths graph kernel

We evaluated the APG kernel using the same cross-validation splits as for the SL kernel. Results shown in Table 3 indicate that experiments achieve similar performance independent of the hyperplane optimization parameter c. Mathematically, a larger generalization parameter c represents a lower risk of overfitting [30, 31].

**Table 3.** APG kernel results on the datasets CPI-DS.

| c | Rec. | Spec. | Prec. | F1 | AUC |
| --- | --- | --- | --- | --- | --- |
| 0.25 | 84.8 | 68.1 | 79.3 | 81.8 | 84.2 |
| 0.50 | 85.3 | 66.5 | 78.7 | 81.8 | 84.2 |
| 1.00 | 84.6 | 66.7 | 78.5 | 81.3 | 83.9 |
| 2.00 | 85.6 | 64.9 | 77.9 | 81.5 | 83.5 |

$c$ is hyperplane optimization parameter. Values in percent: Rec. - recall, Spec. - specificity, Prec. - precision, F1 - F1 score, AUC - area under the curve.

### Both kernels in comparison

In general, the APG kernel achieved slightly better results than the SL kernel in terms of the resulting AUC value, which is inline with previous findings for other domains, e.g. drug-drug interactions [15] and protein-protein interactions [1](Fig 2). Considering the models with the highest precision values for SL (n=3, w=1), and APG (c=0.25) kernel, the results clearly outperform the baseline approach of simple co-occurrences.

**Fig 2.**
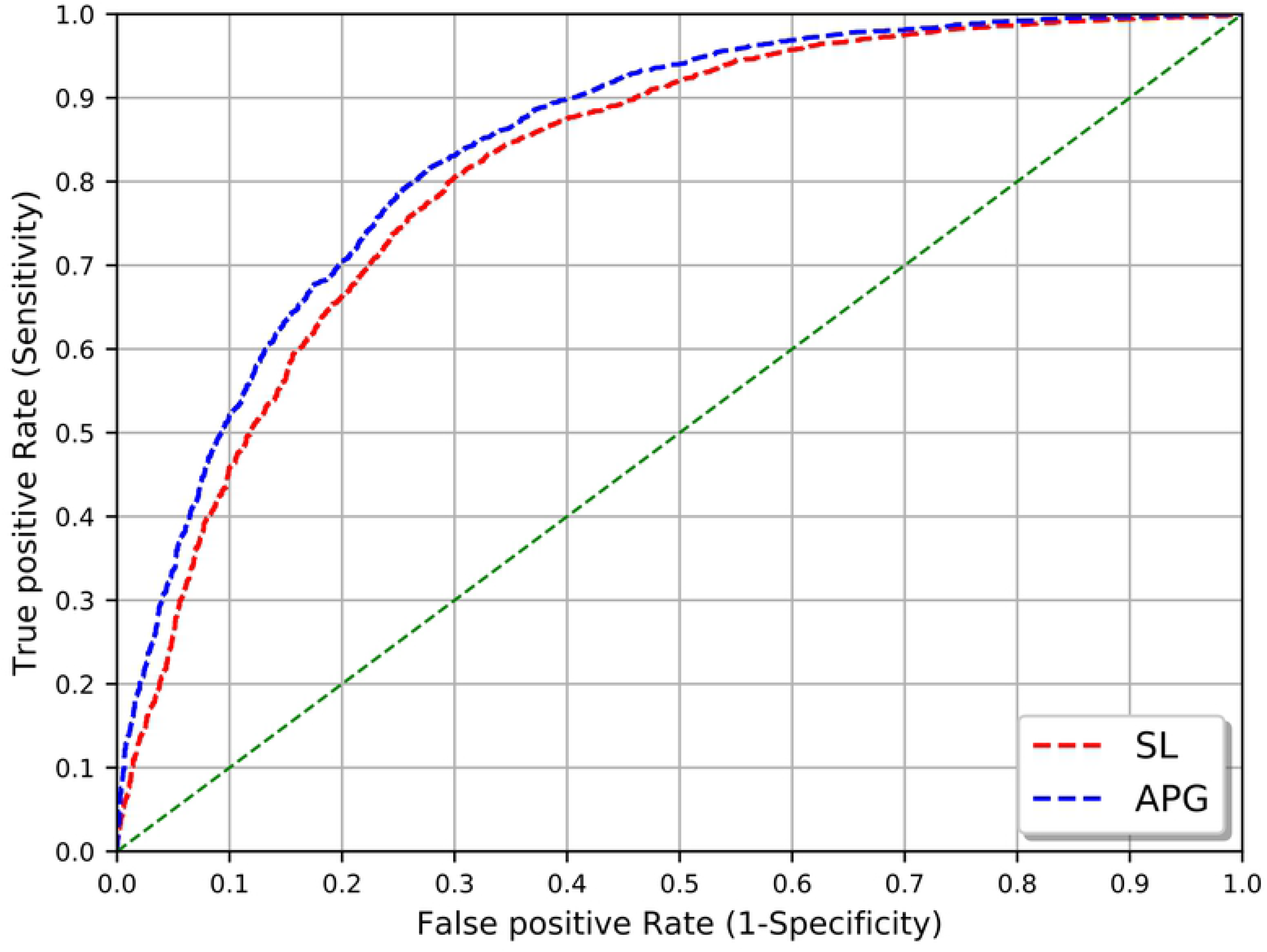
SL and APG kernel comparison. Area under the curve (AUC) of SL kernel (n=3, w=1) and APG kernel (c=0.25).

The complete cross-validation procedure for all parameter settings (including linguistic preprocessing) required almost 5.5 h for the APG kernel and about 35 min for the SL kernel on an Intel Core i5-3570 (4x 3.40GHz). For the APG kernel, a substantial amount of the time is required for dependency parsing. This aspect has to be considered within the scenario of applying a selected model to all PubMed articles (see section *Large-scale dataset application*).

### Both kernels in combination

The combination of both kernels by a jury decision rule yielded a precision of 83.9% and an F1 score of 78.3%. As described in the methods section, the precision was set to the same value (83.9%) for each individual kernel and the appropriate classification threshold was extracted from the AUC analysis. The resulting F1 score was slightly lower for the APG kernel (75.2%) and considerably lower for the SL kernel (70.1%). This indicates that the combination of both kernels by a jury decision leads to a slightly better classification performance(Table 4).

**Table 4.** Comparison of the combined kernels to each individual kernel. The precision of each kernel was set to the same level as in the combination by jury decision.

| Kernel | Rec. | Spec. | Prec. | Acc. | F1 |
| --- | --- | --- | --- | --- | --- |
| SL | 60.2 | 83.7 | **83.9** | 69.9 | 70.1 |
| APG | 68.2 | 81.5 | **83.9** | 73.7 | 75.2 |
| Jury decision | 73.4 | 80.1 | **83.9** | 76.2 | 78.3 |

Values in percent: Rec. - recall, Spec. - specificity, Prec. - precision, F1 - F1 score.

### Functional relationships with and without an enclosed interaction verb

Subsequently, we analyzed the impact of interaction words on the classification of both kernels. The independence of functional relationships and the existence of an interaction verb was tested with a chi-squared test. Both characteristic features are not independent from each other (p<0.01). The fraction of sentences containing an interaction verb is higher in the functionally related CPI-pairs (Fig 3).

**Fig 3.**
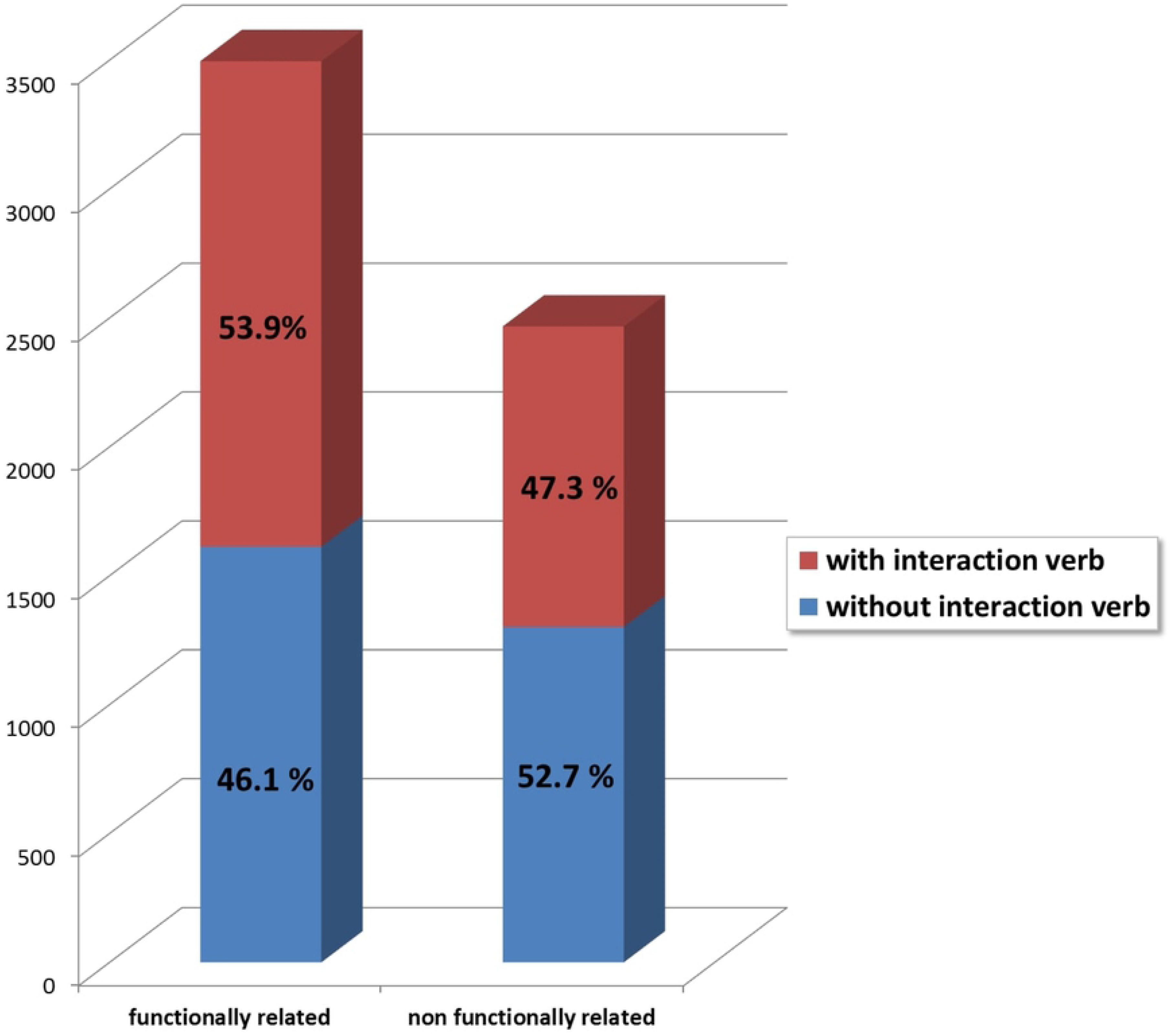
Ratios of CPI-pairs in sentences with and without interaction verbs.

To see if and how the two different kernel functions make use of this correlation, we divided the CPI-DS into two groups, considering compound-protein pairs which contain an interaction verb (CPI-DS_IV) and pairs of compounds and proteins that do not show this structure, i.e. no interaction verb enclosed (CPI-DS_NIV). Table 5 shows the baseline results by using simple co-occurrences. In both datasets, the baseline achieves an F1 score above 70%. Regarding the analyses of the kernels, we recalculated the results from the complete CPI-DS cross-validation run on CPI-DS_IV and CPI-DS_NIV.

**Table 5.**
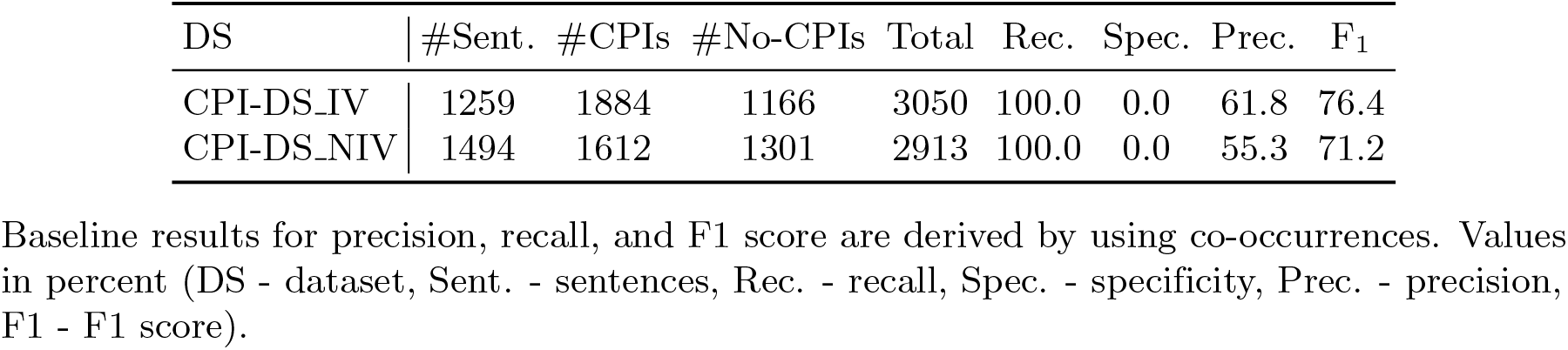
Basic statistic of the two compound-protein interaction corpora.

### Shallow linguistic kernel

For both datasets (CPI-DS_IV and CPI-DS_NIV), the parameter selection n-gram 3 and window size 1 shows the highest area under the curve value (Table 6). Again, a lower value of window size leads to a higher precision and a lower recall. The area under the curve values show the same tendency as the precision, but the differences are not that pronounced. Furthermore, the area under the curve values on the dataset CPI-DS_NIV are about 4-6% higher than on dataset CPI-DS_IV and the specificity about 6-11%. Precision, recall, and F1 score are relatively similar across the two datasets. Therefore, the SL kernel shows a better performance in distinguishing between functional and non-functional relations on dataset CPI-DS_NIV.

**Table 6.** SL kernel results on the datasets CPI-DS_IV and CPI-DS_NIV.

| Param. |  | CPI-DS_IV |  |  |  |  | CPI-DS_NIV |  |  |  |  |
| --- | --- | --- | --- | --- | --- | --- | --- | --- | --- | --- | --- |
| n | w | Rec. | Spec. | Prec. | F1 | AUC | Rec. | Spec. | Prec. | F1 | AUC |
| 1 | 1 | 79.0 | 63.2 | 78.1 | 78.4 | 76.7 | 75.4 | 73.9 | 78.3 | 76.6 | 81.4 |
| 1 | 2 | 84.2 | 57.7 | 76.9 | 80.2 | 76.2 | 81.8 | 65.5 | 74.7 | 77.9 | 80.6 |
| 1 | 3 | 87.6 | 48.4 | 73.7 | 79.8 | 75.8 | 87.4 | 56.4 | 71.5 | 78.5 | 79.9 |
| 2 | 1 | 79.9 | 63.8 | 78.6 | 79.1 | 78.3 | 78.4 | 75.3 | 79.9 | 79.0 | 83.3 |
| 2 | 2 | 83.9 | 57.0 | 76.4 | 79.8 | 77.5 | 84.5 | 67.0 | 76.3 | 80.1 | 82.7 |
| 2 | 3 | 87.7 | 49.7 | 74.2 | 80.2 | 77.1 | 88.2 | 59.4 | 73.1 | 79.8 | 82.3 |
| 3 | 1 | 80.7 | 64.1 | 78.9 | 79.7 | 78.8 | 79.9 | 74.6 | 79.8 | 79.7 | 84.3 |
| 3 | 2 | 83.8 | 58.0 | 76.8 | 80.0 | 78.1 | 84.2 | 68.3 | 76.8 | 80.2 | 83.7 |
| 3 | 3 | 87.7 | 50.0 | 74.4 | 80.3 | 78.0 | 87.9 | 60.8 | 73.6 | 80.0 | 83.2 |

The first parameter shows the n-gram value, and the second number represents the window size. Values in percent: Rec. - recall, Spec. - specificity, Prec. - precision, F1 - F1 score, AUC - area under the curve.

### All-paths graph kernel

Table 7 shows that experiments within the same dataset achieve similar performances, independent of the hyperplane optimization parameter c. For both datasets, the AUC values do not differ by more than 0.8%, indicating a high robustness of the classifier. Furthermore, the AUC values of dataset CPI-DS_NIV are about 3% better than on dataset CPI-DS_IV, due to a clearly higher specificity. Therefore, the APG kernel also shows a better performance in distinguishing between functional and non-functional relations on dataset CPI-DS_NIV.

**Table 7.** CPI-DS_IV and CPI-DS_NIV results for the APG kernel pipeline.

| Param. |  | CPI-DS_IV |  |  |  |  | CPI-DS_NIV |  |  |  |  |
| --- | --- | --- | --- | --- | --- | --- | --- | --- | --- | --- | --- |
| c |  | Rec. | Spec. | Prec. | F1 | AUC | Rec. | Spec. | Prec. | F1 | AUC |
| 0.25 |  | 85.5 | 63.5 | 79.9 | 82.5 | 82.6 | 84.0 | 71.7 | 78.8 | 81.0 | 85.7 |
| 0.50 |  | 85.7 | 61.3 | 79.2 | 82.3 | 82.6 | 84.8 | 70.6 | 78.3 | 81.2 | 85.6 |
| 1.00 |  | 85.2 | 61.0 | 78.9 | 81.8 | 82.3 | 83.9 | 71.1 | 78.4 | 80.7 | 85.3 |
| 2.00 |  | 86.0 | 59.4 | 78.5 | 81.9 | 81.8 | 85.2 | 69.2 | 77.6 | 81.0 | 85.0 |

Values in percent: Rec. - recall, Prec. - precision, F1 - F1 score, AUC - area under the curve.

### Large-scale dataset application

The kernels have been successfully applied to all PubMed titles and abstracts that were published before 2018, comprising about 28M references to biomedical articles. The dataset consists of more than 120M sentences, with around 16M containing at least one compound-protein pair. Table 8 shows that the APG kernel predicts 65.0% of the potential candidate pairs to be functionally related, while the SL kernel predicts only 59.4% as functional. On an Intel Core i5-3570 (4x 3.40GHz), the total run-time of the SL kernel was moderately less than the one of the APG kernel.

**Table 8.** Application of CPI-Pipeline on PubMed dataset.

| Kernel | APG | SL |
| --- | --- | --- |
| PubMed articles | 28M |  |
| Number of sentences | 120M |  |
| Number of sentences with candidate pairs | 5.6M |  |
| Number of candidate pairs | 16M |  |
| Functional relations | 10.4M = 65.0% | 9.5M = 59.4% |
| Non-functional relations | 5.6M = 35.0% | 6.5M = 40.6% |
| Number of identical predictions | 12M = 75%<br>(50% functional, 25% non-functional) |  |
| Number of predicted distinct functional relations | 2.3M |  |
| Pre-processing elapsed time | 20.5 days | 14 days |
| Kernel elapsed time | 4 days | 1.5 days |
| Total elapsed time | 24.5 days | 15.5 days |

PubMed dataset statistics for the selected APG (c=1) and SL (n=3, w=1) models.

## Conclusion

The SL and APG kernels were already applied in different domains, e.g. protein-protein interactions, drug-drug interactions, and neuroanatomical statements. The approach presented herein is focusing on the extraction of functional compound-protein relationships from literature. A benchmark dataset was developed to evaluate both kernels with a range of parameters. This corpus consists of 2,753 sentences, manually annotated with 5,963 compound-protein relationships after automatic named entity recognition. Both kernels are dependent on successfully recognizing protein and compound names in texts. Considering PubMed articles, reliable annotations of these entities are provided by PubTator [33], and publications can be locally processed with PubMedPortable [25].

Cross-validation results with an AUC value of around 81% for the SL kernel and 84% for the APG kernel represent a remarkable performance within the research area of relation extraction.

Considering the filtering of specific interaction verbs, both kernels show the same tendency. The AUC values are considerably higher for the sentences that do not contain interaction verbs. This tendency is not reflected by the F1 score, because this evaluation metric does not include the true negatives. To clarify this outcome, we included the specificity values in our results section. Therefore, the presence of an interaction verb makes it more difficult for the classifier to distinguish between functionally and non-functionally related CPI-pairs. Since the filtering by interaction verbs does not yield a clearly better precision for both datasets and kernels, this approach can be ignored for the development of an automatic classification method. The combination of both kernels could slightly increase the overall performance of the classification compared to the single APG kernel. Since both kernels are quite different regarding their classification approach, their combination is supposed to result in a high robustness of the predictions. For fully automatic methods the classification threshold for both kernels can be adjusted to a relatively high precision. The models for predicting functional relationships between compounds and proteins can then be considered e.g. as a filter to decrease the number of sentences a user has to read for the identification of specific relationships.

The selected procedure of training a model with the SL and APG kernels might also τn come into question for the identification of other types of relationships, such as gene-disease or compound-compound relationships.

## Acknowledgments

We want to thank Kevin Selm for modifying the jSRE software implementation of Giuliano *et al.* to enable parameter selection. We also want to thank Elham Abassian 28: for her support in programming.

